# Construction of a High-Density Genetic Map Based on Large-Scale Marker Development in *Coix lacryma-jobi* L. Using Specifific-Locus Amplifified Fragment Sequencing (SLAF-seq)

**DOI:** 10.1101/2022.02.21.481382

**Authors:** Chenglong Yang, Xiuwen Ban, Mingqiang Zhou, Yu Zhou, Kai Luo, Xiaoyu Yang, Zhifang Li, Fanzhi Liu, Qing Li, Yahong Luo, Xiang Zhou, Jing Lei, Peilin Long, Jian Wang, Jianchun Guo

**Author notes:** Correspondence; (W.J.). These authors contributed equally to this work (Chenglong Yang, Xiuwen Ban, Mingqiang Zhou).

## Abstract

*Coix lacryma-jobi* L. is one of the most important economical and medicine corn. In this study, a high genetic linkage map has been constructed for *Coix lacryma-jobi* L. from a cross F2 community of “Qianyi NO.2” × “Wenyi NO.2” and their parents through high-throughput sequencing and specfic-locus amplified fragment (SLAF) library construction. After pre-processing, 325.49 Gb of raw data containing 1,628 M reads were obtained. A total of 22,944 high-quality SLAFs were detected, among which 3,952 SLAFs and 3,646 of the polymorphic markers could meet the requirements for construction of a genetic linkage map. The integrated map contains 3,605 high quality SLAFs, which are grouped to 10 genetic linkage groups. The total length of the linkage map is 1,620.39 cM with an average distance of 0.45 cM and 360.5 markers in average of per linkage group. This work will provide an important molecular biology basis for investigating their characteristics, gene cloning, molecular marker-assisted breeding, functional genomics and so on for Job’s tears.

## Introduction

*Coix lacryma-jobi* L. (Job’s tears), also named as medicine corn, myotonin, and six grains, is an annual or perennial C4 herb belonging to Coix L. species, Maydeae, Gramineae.. It has been widely grown in East and Southeast Asia[1]. Therefore, southwestern China is one of its centers for the origin, evolution and migration[2,3]. Job’s tears is a traditional crop with high nutritional and one of the most important components of Chinese traditional herbal medicine[4,5,6]. As an application of its seed oil, Kanglaite injection (KLT) has been widely used for cancer therapy[7,8]. With the widespread functional recognition on its nutrition and activities on anti-tumor, immunomodulation, as well as blood-lowering blood calcium, the demand for Job’s tears has increased rapidly and widely as a medicinal and health product in almost all tropical and subtropical countries all over the world[2].

To date, traditional breeding including wild resource domestication, hybrid and mutagenic breeding, has been mainly utilized to breed new varieties of Job’s tears. However, the breeding of Job’s tears was limited due to some of the inherent characteristics and breeding technique limitations. Many important economic traits of the Job’s tears are controlled by multiple gene loci. To improve the important agronomic traits and the breeding efficiency of the Job’s tears, technologies of marker assistive technology (MAS)and marker quantity trait sites (QTLs) can be used to locate and clone important agronomic trait genes. Genetic maps, especially high-density genetic maps, are important tools for QTL location and marker-assisted selection research. Li et al. has used RAPD technology for genetical evaluation on seed resources of Job’s tears[9]. Ma et al. has analyzed the genetic relationship of 79 varieties and found that the genetic diversity of the varieties in Guangxi, China is higher than that in South Korean[10]. Fu et al. has researched on the genetic relationship evaluation with 139 Job’s tears varieties using AFLP and demonstrated that southwestern China is its secondary origin center[3]. Qin et al. has built a genetic map with F2 community of 131 individuals including 10 genetic clusters, 80 AFLP markers and 10 RFLP markers with a total length of 1,339.5 cM, average genetic spacing of 14.88 cM, based on their parents from Beijing and Wuhan[11]. Guo et al. has sketched the whole genome of species “Great Montenegro” with a total genome of 1.6 Gb and mapped the Genetic linkages constructed from 551 individuals of F2 community from “Great Montenegro” (Male parent) x “small white shell Xingren” (Female parent) and BC4 backcrossed by “small white shell Xingren”. This mapcontained 230 Indel markers, a total length of 1,570.12 cM, and average genetic spacing of 6.83 cM. They also accurately identified a gene Ccph1controling the thickness of seed’s shell and *Ccph2* affecting color of seed’s shell[12].

Some genetic maps have been constructed by molecular marking techniques such as AFLP, RFLP, RAPD, ISSR, SRAP, and SSR. Due to the limited amount of individuals used in these maps, low number of molecular markers and the marking density are not saturated enough, which implies difficulty to carry out the follow-up studies such as QTL. Specific site amplification fragment sequencing (SLAF-seq) technology is an effective method for large-volume single nucleotide polymorphism (SNP) and large-scale genotyping based on simplified genomic library (RRL) and high-throughput sequencing[13]. SLAF-seq’s powerful role on genetic research is subsequently used to develop SLAF markers in Lophopyrum elongatum[14] and Corn[15], and high-density genetic mapping construction and QTL location in species such as sesame seeds[16], soybeans[16] and mango[17]. SLAF-seq is the best choice for large-scale molecular marker development and high-density chain mapping, especially in organisms that do not have reference to genomic information.

After several years of investigation and observation, two personality traits with large differences were chosen for hybridizing and their genetic linkage map. A large number of SNP markers have been identified by using SLAF-seq method, and these new markers were used to construct a high-density genetic linkage map, which could provide theoretical basis for investigating their characteristics, gene cloning, molecular marker-assisted breeding, functional genomics and so on.

## Materials and Methods

### Plant Materials and DNA Extraction

200 individuals were randomly selected from F2 separation group of 426 individuals which were built from “Wenyi NO.2” (male parent) x “Qianyi NO.2” (female parent) as genetic mapping separation group. These two parents and F2 generations were planted at the Institute of Subtropical Crops of the Guizhou Academy of Agricultural Sciences. The healthy leaves from parents and F2 generation were collected and stored in liquid nitrogen. For DNA extraction, the CTAB buffer (8.18 g NaCl, 2 g CTAB, in total volume of 100 mL with 20 mM EDTA, 100 mM Tris, pH 8.0) has been modified based on the traditional CTAB method[18]. The total genomic DNAs were respectively extracted from each plant and analyzed by electrophoresis with 1% agarose gel and quantified by spectrophotometer [19](NanoDrop 2000, Thermo, USA).

### SLAF library Construction and High-throughput Sequencing

SLAF-Seq method was used[20]. Firstly, Genome of the parents and F2 populations was digested by Hpy166II restrictive endoenzyme (New England Biolabs (NEB), USA); then the single nucleotide (A) was added to the end of the digestive fragment by Klenow fragment (3’-5’ exo-) (NEB) with dATP at 37°C. Afterwards, the dual-label sequencing markers (PAGE-purified, Life Technologies, USA) were connected to the new added terminal A by T4-DNA connecting enzymes. PCR amplification was performed with the diluted DNA samples, primers of 5’-AATGATACCGACCACCGA-3’ (forward) and 5’-CAAGCAGAAGACGGCATA-3’(reverse), Q5^®^ High-Fidelity DNA Polymerase (NEB), and dNTPs. The PCR products were purified and collected by Agencourt AMPure XP beads (Beckman Coulter, High Wycombe, UK) and separated by electrophoresis in 2% agar gel. The DNA fragment (with indices and adaptors) between 264 bp and 464 bp were re-separated and purified from the gel band in electrophoresis by QIA quick gel extraction kit (Qiagen, Hilden Qiagen, Germany); and the paired obtained sequences (terminal 125 bp) were analyzed by Illumina Hi-Seq 2500 system (Illumina, Inc., San Diego, CA, USA).

### SLAF-Seq Data and Genotyping Analyses

SLAF-seq data analysis and genotyping method of Sun[20] were used. Dual index[21] was selected for raw data sequence identification, reads for each sample were used to evaluate the quality and quantity of the sequencing data. SLAF labels were also developed in the parent and F2 community through reads clustering. And polymorphism analysis was performed based on the difference between the number of allelic genes and the gene sequences. The SLAF labels (polymorphic SLAF label) with polymorphic sites (SNP and Indel) were selected for subsequent analysis. The SLAF markers were evaluated and filtered multiple times to obtain the high-quality, effective molecular markers. Since the Job’s tears is a diploid plant, the filtered DNAs contained more than 4 different suspicious SLAF genotypes at one gene site. In this study, the sequence length less than 200 bp was defined as a low length of SLAFs and filtered out. SLAFs with 2, 3, or 4 tags were identified as polymorphic SLAFs and considered as potential tags. Polymorphic markers were divided into 8 separation modes, which were ab×cd, ef×eg, hk×hk, lm×ll, nn×np, aa×bb, ab×cc and cc×ab. The F2 population was obtained through F1, a fully pure parent with two genotypes of aa or bb. Therefore, the SLAF marker of separation pattern aa×bb was used for genetic map construction.

### Genetic Map Construction

Linkage groups (LGs) have been initially divided by improved LOG score (MLOD) value up to 5 on mark site. To build more effectively maps, High Map strategy was chosen for arranging SLAF tags in a specific order and correcting genotyping errors in LGs[22]. The genetic map was constructed by maximum likelihood method[23], and genotyping errors will be corrected by smooth method[24]. The missed genotype will be estimated by k-nearest neighbor algorithm[25]. The recombinant r value will be converted to genetic map distance through Kosanbi function (centimorgan, cM)[26] and a high-density genetic chain map of Job’s tears has been drawn. Areas with 3 or more than 3 partial separation markers in adjacent locations on map become the area of the partial separation hot spot which were defined as Segregation distortions region (SDR)[27].

## Results

### Analysis of SLAF-seq Data and SLAF markers

Based on SLAF-Seq technology, total of 1,628,398,591 reads with approximately 325.49 Gb of raw data (Table S1) have been obtained by two-terminal sequencing on the constructed SLAF library. The effective length of each read is approximately between 384 bp to 464 bp by removing the label sequence at the end of the DNA fragments. The average Q30 of the high-throughput sequencing results is about 93.92% and the GC content ranges from 44.43% to 49.75% with average of 46.96% in the 200 F2 community. In order to improve the efficiency of molecular markers, the length of SLAF sequencing for the parent is much greater than the F2 generation. Therefore, in the total reads, 66,927,932 reads originate from the male parent, and 52,144,104 reads originate from the female parent. And in the 200 F2 generation, the sample sequences range from 2,878,250 to 14,176,926, with an average value of 7,546,632 (Table S1).

### SLAF Mark detection and Genotype Definition

All sequences are formed SLAF clusters according to similarity clustering. After eliminating sequences with low lengths, repeats, and suspicious SLAF, the valid SLAF marks are 262,222 and 219,948 in the male and female parents, respectively through high-throughput sequencing. The developed SLAF marks are18,747,434 and 13,716,859 in the male and female parents, of which the average developed rates are 71.50 and 62.37 folds, respectively. In the F2 community, the SLAF marks range from 110,940 to 197,542 with an average of 153,652 through high-throughput sequencing respectively. The developed SLAF marks range from 639,392 to 4,736,985 with an average of 1,969,377, of which the average developed rates are from 5.19 to 30.27 × with total average coverage rate of 12.82 for per individual (Figure 1).

**Figure 1.**
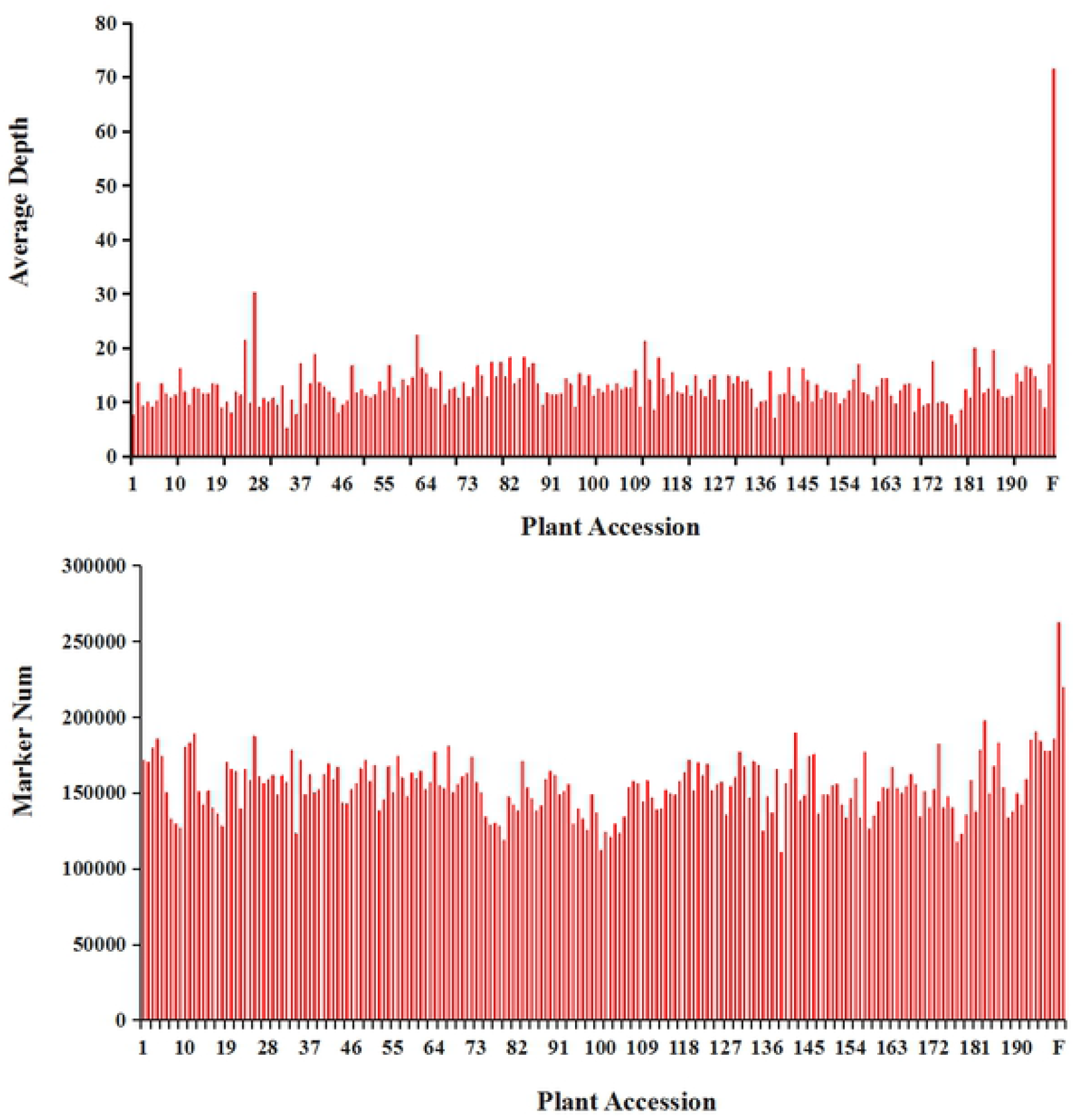
Number and coverage of valid SLAF markers for each F2 individual. The x-axis indicates the plant accession including female parent (F), male parent (M), and each of the F2 individuals; the y-axis indicates the number of SLAF markers (A) and the coverage (B).

Based on the polymorphism differences between alleles and gene sequences, 302,295 high-quality SLAF markers have been developed to 3 types, which are polymorphic markers, non-polymorphic markers and repetitive polymorphism slotted markers (Table 1). Among them 79,364 markers belong to polymorphic markers (26.25%), which are used as subsequent genotypes. After removing low-quality polymorphism markers such as missing parent information, repeat sequence areas, and low integrity of the non-polymorphic markers (73.08%) and repeat polymorphic markers (0.67%), 53,023 of high-quality and effective SLAF markers are genetically classified into 8 separation mods, which are ab×cd, ef×eg, hk×hk, lm×ll, nn×np, aa×bb, ab×cc, and cc×ab (Figure 2). The generation of F2 community are constructed by self-inbred of the homozygote aa with bb hybridize. Therefore, only aa×bb isolates are used for genetic map construction and 22,944 SLAF markers are genetically classified into this group, and 3,952 of them originate from the parents with average sequencing depth of 66.925×; and in the F2 generation, the sequencing depth is 12.82×. All these SLAF marks were used to genetic maps construction.

**Table 1.** SLAF mining results.

| Type | Number of SLAF markers | Number of reads | Ratio |
| --- | --- | --- | --- |
| Polymorphism | 79,364 | 110,270,167 | 26.25% |
| Non-polymorphism | 220,904 | 311,617,085 | 73.08% |
| Repetitive | 2,027 | 2,895,714 | 0.67% |
| Total | 302,295 | 422,400,985 | 100% |

**Figure 2.**
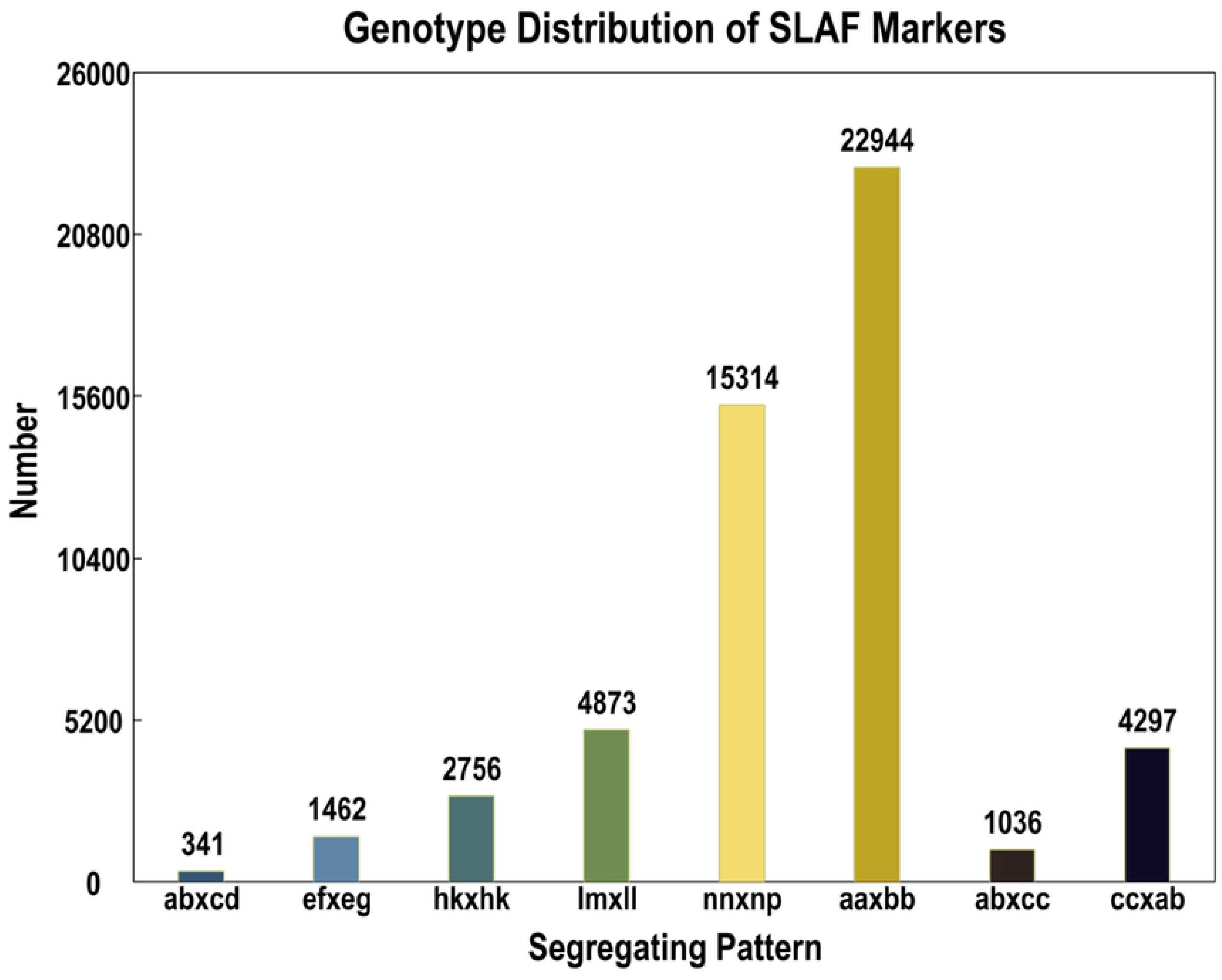
Number of polymorphic SLAF markers for eight segregation patterns. The x-axis indicates eight segregation patterns of polymorphic SLAF markers; the y-axis indicates the number of markers.

### Basic Characteristics of the Genetic Map

After a series of screenings, 3,605 effective SLAF markers from total of 3,646 ones were obtained and used for the final linkage analysis, while the rest 41 SLAF markers were not located on any of the linkage map. These markers have 165.30 × and 98.59 × of coverage in the male and female parents, respectively; and 20.14 × of the average coverage in the F2 individuals. In 200 of F2 individuals, the integrity of each marker is a key parameter in controlling the quality of the genetic map. The averages integrity of all markers positioned on chain map reaches up to 98.95%.

3,952 SLAF markers from 202 individuals has been used for genetic mapping. Linkage analysis through Mendel separation ratio was also achieved by software of Joimnap 4.1 in conditions of LOD ≥ 4.0 and recombination rate (r) ≤ 0.30,3,605 SLAF markers from total of 3,646 have been distributed across 10 linkage groups. The present rate reaches up to 98.88%. Finally, analysis of the distribution of all markers on 10 linkage groups showed that the total genetic distance was 1,620.39 cM and the average distance between two SLAF markers was 0.45 cM, which is the densest genetic linkage map as far as in Job’s tears. The marker distribution and length of the 10 linkage groups were not the same (Table 2 and Figure 3). The largest LG is LG9 with 774 markers with the total length of 266.78 cM, and the average distance between adjacent markers is only 0.35 cM; the smallest LG is LG6 with only 97 markers with a length of 66.73 cM, and the average length between adjacent markers is only 0.70 cM. In the 10 linkage groups, the average SLAF markers are 360.5, the genetic LG length ranges from 66.73 cM (LG6) to 266.78 cM (LG9), and the distance between adjacent markers are between 0.35 cM (LG9) and 0.84 cM (LG10), the linkage levels range from 98.06% to 100.00% with an average of 99. 42% in conditions of “Gap≦5”, the largest interval length is 9.72 cM (LG5) (Table S2). In the above LGs, 10 segregation distortion regions that are 0.28% of the total SLAF markers are found, and 8 of them deviate in the male-parents and 2 of them deviate in the female-parents. 8 of the 10 segregation distortion regions (SDRs) locate in LG7, and the remaining 2 distribute in LG9.

**Table 2.** Distribution of SNP loci on the 10 linkage groups of coix.

| LG ID | Marker number | Total Distance (cM) | Average Distance (cM) | Gaps<=5(%)<sup>1</sup> | Max Gap | SDR |
| --- | --- | --- | --- | --- | --- | --- |
| LG1 | 428 | 157.79 | 0.37 | 100 | 3.71 | 0 |
| LG2 | 229 | 155.38 | 0.68 | 99.56 | 8.36 | 0 |
| LG3 | 336 | 175.84 | 0.52 | 99.1 | 6.82 | 0 |
| LG4 | 412 | 172.84 | 0.42 | 99.76 | 5.87 | 0 |
| LG5 | 411 | 157.82 | 0.38 | 99.51 | 9.72 | 0 |
| LG6 | 97 | 66.73 | 0.7 | 100 | 2.83 | 0 |
| LG7 | 459 | 183.62 | 0.4 | 99.56 | 7.15 | 8 |
| LG8 | 303 | 153.86 | 0.51 | 98.68 | 9.02 | 0 |
| LG9 | 774 | 266.78 | 0.35 | 100 | 4.67 | 2 |
| LG10 | 156 | 129.73 | 0.84 | 98.06 | 9.32 | 0 |
| Maximum | 774 | 266.78 | 0.84 | 100 | 9.72 | 8 |
| Minmum | 97 | 66.73 | 0.35 | 98.06 | 3.71 | 0 |
| Total | 3,605 | 1,620.39 | / | / | / | 10 |
| Average | 360.5 | 162.04 | 0.45 | 99.42 | 9.72 | / |

<sup>1</sup>Gap < =5' indicated the percentages of gaps in which the distance between adjacent markers was smaller than 5 cM.

**Figure 3.**
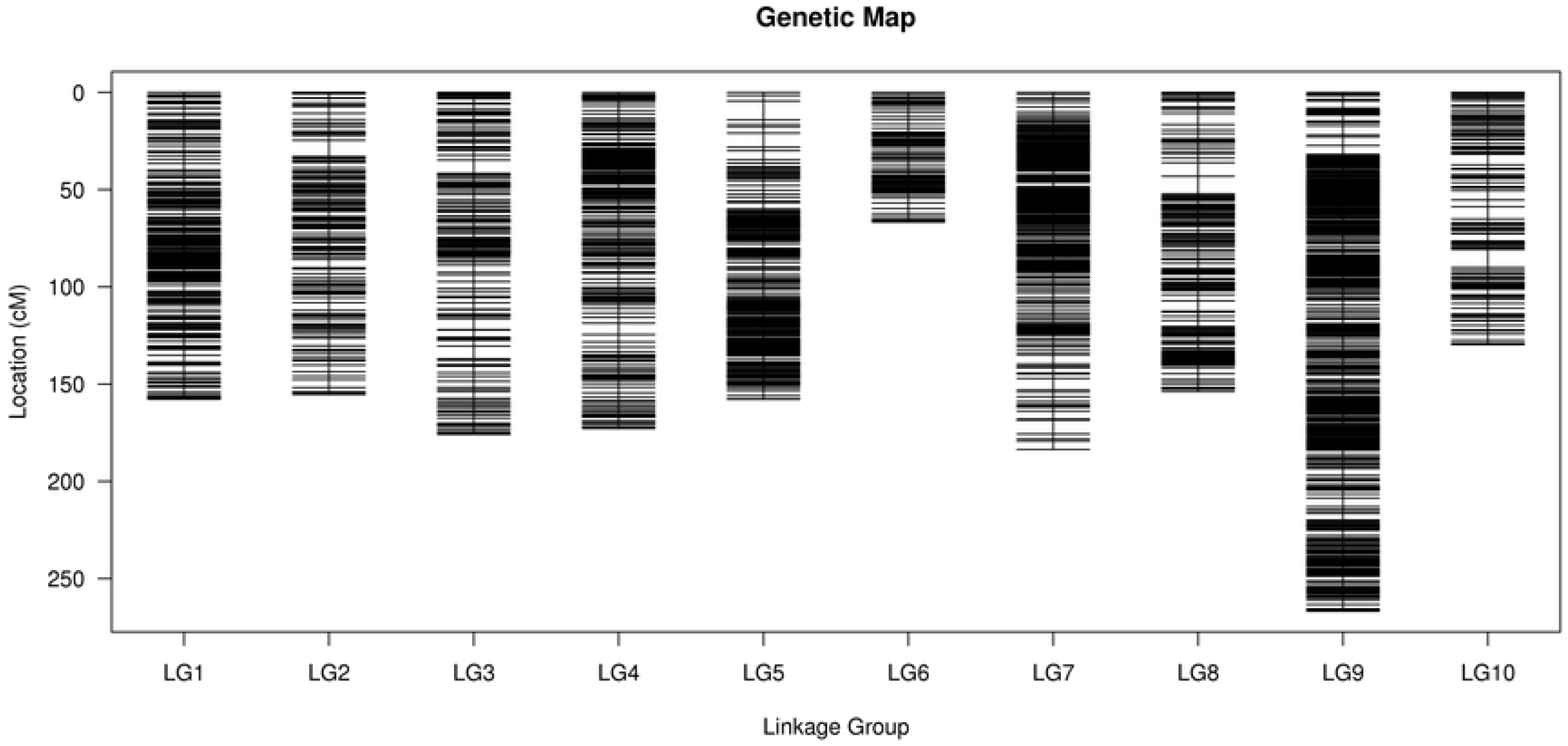
Distribution of SLAF markers on the 10 linkage groups of Job’s tears. A black bar indicates a SLAF marker. A red bar indicates a segregation distortion marker. The x-axis represents the linkage group number and the y-axis indicates the genetic distance (cM) in each linkage group.

### Distribution of Markers on the Genetic Map

Total of 5,378 SNP markers, 1,519 SNP switch points 3, 859 SNP conversion points have been obtained in the constructed genetic map with 3,605 SLAF markers, and the ratio between SNP switch and SNP conversion points is 0.39. The distribution rule of the markers has also been analyzed for each LG of the 10 LGs. The results showed that the largest quantity of SNP markers are 1,382 in LG9, and 372 of them belong to SNP switch points, the rest 1,010 belong to SNP conversion points (Table 3). The ratio of SNP switch to SNP conversion points is 0.37. LG6 has the smallest number of 147 SNP markers, and 54 of them belong to SNP switch points, the rest 93 belong to SNP conversion points, and the ratio of SNP switch and SNP conversion points is 0.58 (Table 3). Total of 5,387 SNP loci are detected among the 3,605 SLAF markers on the final map (Table 3). Further analysis has been executed on these SNPs, which show that the transition type is majority SNPs, and reachs up to 71.76%, including the R (A/G) and Y types (C/T) with 35.35% and 36.41% respectively. The transversion type is less with 28.4%, including M (A/C), W (A/T), S (C/G) and K(G/T) types, among which the ratios range from 6.66% to 7.49% respectively (Table 4).

**Table 3.** Statistic of mapped SNP marker types.

| Linkage group | SNP number | Transition/Transversion number |
| --- | --- | --- |
| LG1 | 541 | 388/153 |
| LG2 | 268 | 194/74 |
| LG3 | 440 | 320/120 |
| LG4 | 623 | 452/171 |
| LG5 | 613 | 418/195 |
| LG6 | 147 | 93/54 |
| LG7 | 755 | 545/210 |
| LG8 | 428 | 308/120 |
| LG9 | 1382 | 1,010/372 |
| LG10 | 181 | 131/50 |

**Table 4.** Statistic of mapped SNP marker types

| SNP types | Number | Ratio (%) |
| --- | --- | --- |
| R(G/A) | 1,901 | 35.35 |
| Y(T/C) | 1,958 | 36.41 |
| S(G/C) | 392 | 7.29 |
| W(A/T) | 403 | 7.49 |
| K(G/T) | 358 | 6.66 |
| M(A/C) | 366 | 6.81 |
| Total | 5,378 | 100 |

### Quality evaluation of Genetic map

The quality genetic map of Job’s tears has been evaluated through haploid and thermal map construction. The haploid map reflects the double exchange from the parents and the 200 F2 generations by 3,605 SLAF markers (Supplementary Material Presentation 1). Most recombinant areas can be clearly determined, and color changes indicate the occurrence of reorganization events. A large proportion of the recombinant regions are determined by their color differences. The missing markers for each LG range from 0.05% (LG4) to 1.9% (LG7). Most LGs are evenly distributed, which showed the genetic map has very high quality (Table 5).

**Table 5.** Distribution of segregation distortion markers.

| Linkage<br>group | All marker | | Segregation distortion marker | | $\chi^2$ | *P* | Frequency of segregation<br>distortion marker | SDR number |
| --- | --- | --- | --- | --- | --- | --- | --- | --- |
|  | Number | Percentage | Number | Percentage |  |  |  |  |
| 1 | 428 | 11.87% | 88 | 14.84% | 0.465 | 0.203 | 20.56% | 0 |
| 2 | 229 | 6.35% | 20 | 3.37% | 0.865 | 0.862 | 8.73% | 0 |
| 3 | 336 | 9.32% | 6 | 1.01% | 0.008 | 0.158 | 1.79% | 0 |
| 4 | 412 | 11.43% | 68 | 11.47% | 1.213 | 0.032 | 16.50% | 0 |
| 5 | 411 | 11.40% | 17 | 2.87% | 0.043 | 0.313 | 4.14% | 0 |
| 6 | 97 | 2.69% | 15 | 2.53% | 0.527 | 0.005 | 15.46% | 0 |
| 7 | 459 | 12.73% | 127 | 21.42% | 0.628 | 0.014 | 27.67% | 8 |
| 8 | 303 | 8.40% | 99 | 16.69% | 0.802 | 0.307 | 32.67% | 0 |
| 9 | 774 | 21.47% | 101 | 17.03% | 0.032 | 0.504 | 13.05% | 2 |
| 10 | 156 | 4.33% | 52 | 8.77% | 0.824 | 0.014 | 33.33% | 0 |
| Total | 3,605 |  | 593 |  |  |  | 17.39% | 10 |

$\chi^2$ and *P* indicate c2 values with one degree of freedom and the corresponding probability, respectively. SDR means segregation distortion region.

The genetic map is a basic multi-point recombination analysis, of which the closer distance between markers, the smaller recombination rate, the potential layout problems could be found through recombination relationship analysis between Markers and the surrounding Markers. The recombination relationship between LG markers can be demonstrated by thermography map, which has been constructed from the multiple comparisons of the recombination levels from the SLAF 3,605 markers (Supplementary Material Presentation 2). The linkage relationship between adjacent markers on each linkage group is addtion with the increase of distance, the linkage relationship between the markers and distant markers is gradually weaker indicating that marker sequences are correct.

### Segregation Distortion Markers on the Map

The chi-square test showed 593 out of total 3,605 SLAF markers on genetic map are distortion markers which reached up to 16.45% under P<0.05. The distortion markers are distributed on all 10 linkage groups with different proportions, and their distribution mode in each LG is similar to all marker distribution, except in LG7, LG8 and LG10 (Table 5). The LG10 has the highest percentage of distortion markers which reach up to 33.33%. The LG3 has the lowest percentage of distortion markers with 1.79%. The largest LG9 has 101 distortion markers with a distortion ratio of 13.05%. In LG8 and LG10 the frequently of distortion markers are significantly higher than other LGs which are 32.67% and 33.33% respectively.

## Discussion

### The Feasibility and Advantages of the SLAF-Seq Technique

Developing abundant and reliable molecular marker is of great significance for genetic map construction. And SNP marker has been used for genetic map construction in this research. Comparing to the common method, SLAF-seq technology used for large-scale marker development provided a higher density, better consistency, better effectiveness and saved the cost[20]. SLAF technology has been used in a large amount of plant molecular marker development and genetic mapping for laurel[28], mango[29], duck [30], soybeans[31] and other important crops. The smooth development of this study proved that SLAF sequencing technology is an effective method for large-scale development of molecular markers and high-density genetic mapping.

In this study, SLAF reduced-representation genome sequencing method has been used for large-scale marker development of Job’s tear. The high-throughput sequencing of Job’s tears SLAF library has been achieved under Illumina sequencing platform leading to a raw data of 325.49Gb for total of 1,628,398,591 double-ended sequencing fragments. Total of 302,295 SLAF markers were obtained by comparing and clustering analysis, and 3,605 effective markers with high polymorphism were received after filtering and eliminating the low-quality markers which was used for Job’s tears genetic linkage map construction. In this study, large number of molecular markers of Job’s tears were constructed to supply an important foundation for genetic map construction. Abundant of genome information provided an important reference for further molecular biology investigation on Job’s tears.

An important advantage of SLAF sequencing technology is the development of large-scale molecular markers in a single experiment. However, the sequence data obtained by high-throughput sequencing inevitably included numerical missing and errors. Therefore, multi-selection must be used for eliminating the errors which can affect genetic map construction[20]. 79,364 polymorphic SLAF markers has been obtained from the raw sequence data in this study but only 3,605 effective SALF markers has been final received after eliminating the markers possibly due to parent-genotype missing, low integrity, Mendelian error and significant segregation. In addition, sequencing result indicated that the accuracy of SLAF can be effectively improved by increasing the sequencing depth which is consistent with previous studies[20,32]. Therefore, it is necessary to improve the depth of sequencing in future experiments for improving the efficiency of marker developing.

In this study the GC content of Job’s tears SLAF library was 46.96% which is slightly lower than transcription group[33]. This might be due to the different DNA sequence origins (i.e. genomic DNA, cDNA or EST). The types of SNP site in SLAF markers showed that most of them belong to conversion types (71.76%) which was similar to mango, peonies and sesame seeds [34]. The polymorphism rate of SLAF markers among parents of Job’s tears in target group was 26.25% which was lower than EST-SSR polymorphism rate of 31.1%[33]. However, this polymorphic rate of SLAFs is higher than many other varieties[11,12], indicating a high genetic diversity has been constructed among Job’s tears varieties through SLAF which is consistent to previous investigation[2,35].

### Characteristics of the Genetic Linkage Map

By comparison with the other annual herb of homozygous genome, the heterozygosity of Job’s tears genome is far more complex for the genetic map construction. Therefore, Job’s tear has a more complex marker segregation type which is different from other traditional segregation population. In this study, “Wenyi NO.2” and “Qianyi NO.2” were used as male and female parent, respectively, and the F2 community has been used for genetic map. Besides, the Job’s tears has higher heterozygosity with complexed genome reached up to 1.6 to 1.73 Gb approximately, which makes it harder for high-density genetic map construction[12,36]. Therefore, it is particularly important to find proper method for high-density linkage map construction of Job’s tear.

Two-step map that optimizes high-density linkage map construction method has been used for accurating high-density genetic map of Job’s tears construction through κ-Nearest neighbor, maximum likelihood algorithm by repeatedly sorted and error correction strategies[22]. Compared with the traditional JoinMap software, Two-step method is more efficient for high-throughput sequencing, more accurate and reliable for linkage maps construction, more efficient for mapping. This mapping strategy has been widely used in the studies of genetic maps such as Laurel[28], Mango[29], Duck Apotheph[30], and peony[37]. And the powerful power of two-step mapping has been further confirmed in this study for high-density lingage maps construction of Job’s tears.

The first high-density genetic map of Job’s tears has been constructed in our study, and 3,605 SLAF markers cluster to 10 genetic linkage groups were obtained withsame nucleus (2n=20) as the Beijing Job’s tear species[38]. The total length of the map is 1,620.39 cM, with an average genetic distance of 0.45 cM, and 360.5 markers in average of per linkage group. The number of markers on each linkage group are widely variety, some of them are highly clustered in certain region, especially in LG1, LG7 and LG9. This phenomenon migth be caused by factors such as inconsistent molecular marker polymorphism or inconsistent recombination rates in certain chromosome regions of the graph-based parents[13]. Similar phenomena could also be found in plants such aspeony[37], sunflowers[39]. Ma et al. considered that cluster aggregation of molecules in the linkage group is related to the chromosomal filament or hesochtine region[40].

### Segregation Distortion Markers Analysis

Segregation distortion is widespread in nature, which is considered to be one of the important drivers of species evolution, and reasons of segregation distortion are still doubts and controversy[41]. Faure et al. considers that the segregation distortion is formed due to biological factors including the selection of gamete and zygote, non-homogenous recombination, translocation on chromosomes or non-homogenous sites, low homozygosity of parents for mapping[42]. Previous studies have found that environmental factors, experimental errors, the ability of offspring separation and the loss of chromosomes might also be the reason for marker segregation distortion[37,43,44,45,]. In this study, about 16.45% of 593 markers showed significant segregation distortion (P<0.05) in the target group. The high segregation distortion rate further indicated that the genetic diversity of their parents is high. And all 593 segregation distortion markers form 10 SDRs, and cluster distributed in LGs. The aggregation of segregation distortion might be due to the selection of gametophyte and sporophyte[46]. And this phenomenon is widely found in plants[13,16,45]. In addition, the use of segregation distortion markers to build a linkage map can increase the genomic coverage of the genetic map[16,43], and may be conducive to quantitative site (QTL) positioning[47,48].

High-density genetic linkage map has been constructed in this study which has been provided an effective way for important character analysis through QTL, Map-based cloning, and molecular Marker assisted selection on Job’s tear breeding. The 3,605 SNP markers about 93.78% in total map are the common dominant gene sequence labels which could give helps in comparative genomic researches[49] and association mapping[50]. And more important is that the molecular markers of high-density genetic map through SLAF is developed on the genome-wide level.

This study has demonstrated that the application of SLAF-seq technology on large-scale genetic markers of Job’s tears, and HighMap is an effective tool on molecular marker developing and high-density linkage map constructing by using the high-throughput sequencing data. This work provided an important molecular biology basis for genetic diversity, relationship, variety identification, QTL location of phenotypic matters, and genome function structure analysis of Job’s tears.

## Conclusions

We report here the first high-density genetic map for Job’s tears. The map was constructed using an F2 population and the SLAF-seq approach, which allowed the efficient development of a large number of polymorphic markers in a short time. this study may facilitate genomic structure and function, maker-assisted selection breeding, and comparative genomic analyses between Job’s tears and relative species, may further provide an important reference for molecular marker polymerization breeding of Job’s tears, and pave the way for the study of genetic mechanism of important medicinal traits and the origin of Job’s tears.

## Supporting information

**S1 Fig. Haplotype map of the integrated maps**. Each row represents a marker. Markers are ranked in accordance with the map order. Each of the two columns represents an individual; blank columns are used between two individuals. The fifirst and second columns represent the paternal and maternal chromosomes, respectively. The green and blue areas in the columns represent the fifirst and second alleles from the parents, respectively.

The white column represents the source of alleles that cannot be judged. The gray areas represent the deleted alleles.

**S2 Fig. Heat map of the integrated maps**.

Markers of each row and column are ranked according to the map order; each small square represents the rate of recombination (r) between the two markers.

**S1 Table. Summary of the sequencing data. (XLSX)**

**S2 Table. SLAF markers and SNP loci on the map**. The SNPs within the SLAF marker allele sequences were marked in lowercase letters. The SLAF markers separated by linkage groups are shown in different sheets and the parent from which the alleles were derived are shown. (XLSX)

## Author Contributions

The listed authors contributed to this work as described in the following: Conceptualization, CY, MZ and JG gave the concepts of this work, designed the whole experiments, and provided financial support; CY did the experiments and prepared the original draft; CY, JW and XB did part of the experiment and data creation; YZ, KL, PL, XY and ZL reviewed and edited this manuscript; FL, QL, YL, XZ and JL did the methodology and formal analysis. All authors have read and agreed to the published version of the manuscript.

## Acknowledgments

This research was funded by the Natural Science Foundation of China (No. 31760423),Guizhou Agricultural Research Project (No.NY[2015]3020-2, [2017]2508), Guizhou Academy of Agricultural Sciences Special Project (No. [2016]019), Guizhou Social Research Project (No. [2016]2840), Modern agricultural industrial technology system of characteristic coarse grains in Guizhou Province (No. GZCYTX2019-2022).

## Conflicts of Interest

The authors declare no conflict of interest.

## Institutional Review Board Statement

This research does not involve ethics.

## Notes

### Competing Interest Statement

The authors have declared no competing interest.

